# Systemic coagulopathy drives host lethality in a new Drosophila tumor model

**DOI:** 10.1101/2022.05.11.491522

**Authors:** Tsai-Ching Hsi, Katy L Ong, Jorian J Sepers, Jung Kim, David Bilder

## Abstract

Malignant tumors trigger a complex network of inflammatory and wound repair responses, prompting Dvorak’s characterization of tumors as ‘wounds that never heal’ ^1^. Some of these responses lead to profound defects in blood clotting, such as Disseminated Intravascular Coagulopathy (DIC), which correlate with poor prognoses ^2–4^. Here, we demonstrate that a new tumor model in *Drosophila* provokes phenotypes that recapitulate coagulopathies observed in patients. Fly ovarian tumors overproduce multiple secreted components of the clotting cascade and trigger hypercoagulation of fly blood (hemolymph). Hypercoagulation occurs shortly after tumor induction and is transient; it is followed by a hypocoagulative state that is defective in wound healing. Cellular clotting regulators accumulate on the tumor over time and are depleted from the body, suggesting that hypocoagulation is caused by malignant growth exhaustion of host clotting components. Interestingly, clinical studies have suggested that lethality in patients with high serum levels of clotting components can be independent of thrombotic events ^5,6^. We show that rescuing coagulopathy improves survival of tumor-bearing flies, despite the fact that flies have an open circulatory system. Our work establishes a platform for identifying alternative mechanisms by which tumor-driven coagulopathy triggers early mortality, as well as exploring other conserved mechanisms of host responses to chronic wounds.

## RESULTS AND DISCUSSION

### Generation of an ovarian carcinoma model to investigate tumor-host interactions

Tumor-host interactions, as well as autonomous growth of the tumor itself, play central roles in cancer progression, morbidity and mortality ^7,8^. Drosophila has recently emerged as a valuable system to study these interactions, elucidating mechanisms that can be conserved with mammals ^9,10^. However, current systems have key limitations. In larval models, tumor-associated pupation defects prevent lifespan analysis, whereas in adult allograft models, wounding from transplantation confounds the study of the response to the tumor alone. In adult transgenic gut tumor models, perturbation of this essential organ may directly impact host metabolism, physiology, and the microbiome. We therefore developed an alternative genetic model, wherein a tumor is induced to grow in the non-essential ^11^ ovarian epithelium of an adult female fly (**Figure 1A**). In the ovarian carcinoma (OC) model, tumorigenesis is driven by expression of the oncogenes *Ras*^*V12*^ and *aPKC*^*ΔN* 12^, directed to the follicle epithelium via *traffic jam-Gal4 (tj-GAL4)* and restricted to adulthood via a ubiquitously expressed temperature-sensitive GAL4 repressor (*tubGAL80ts)*. This spatiotemporal control facilitates the study of both tumor and adult host from the initial stages of transformation to a fully formed malignancy.

**Figure 1.**
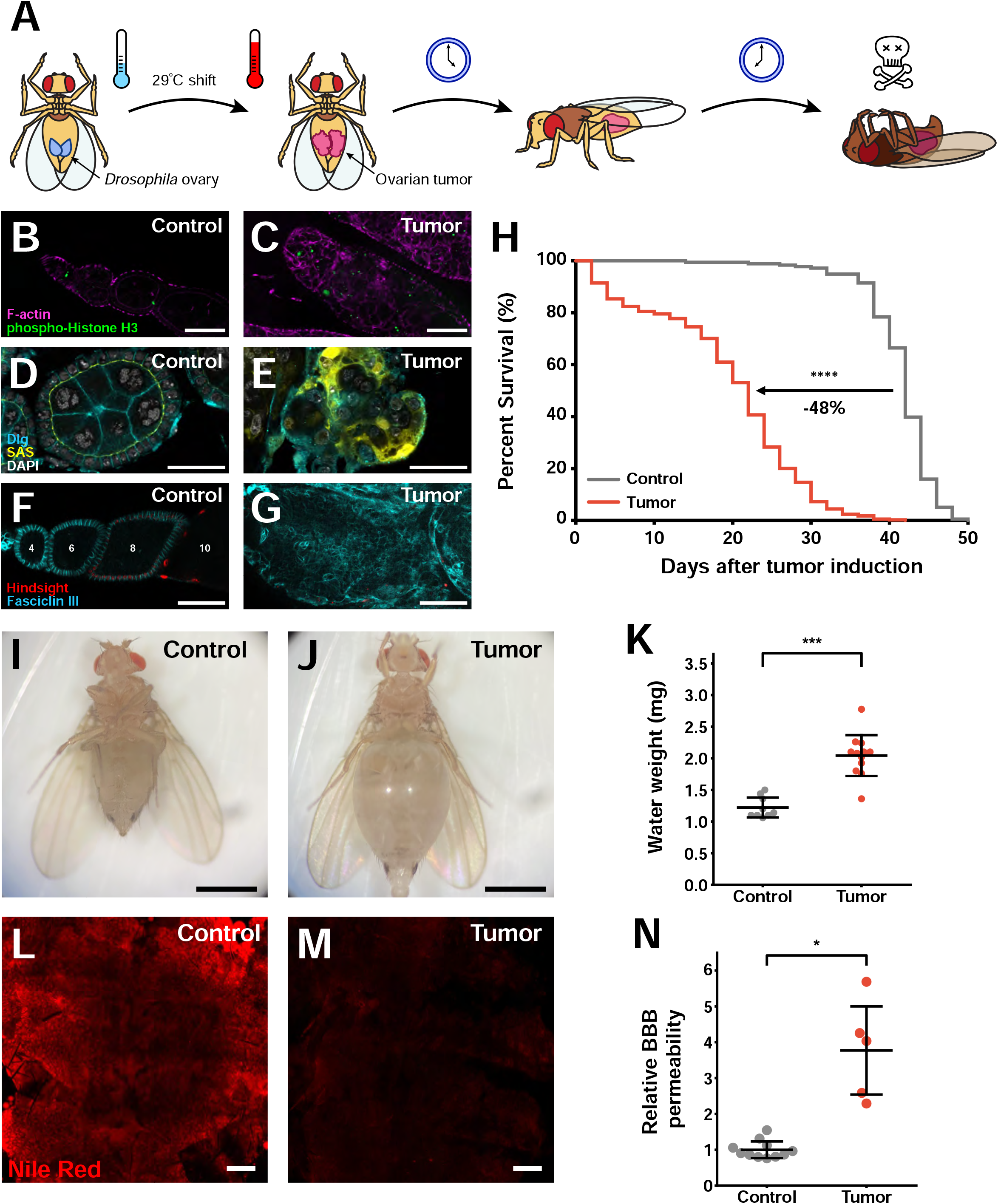
A novel ovarian carcinoma model to study paraneoplasias in *Drosophila*. (A) Schematic of *Drosophila* ovarian carcinoma (OC) induction in adult flies. (B, C) OC tumors exhibit increased anti-phospho-histone H3 staining (green). (D, E) SAS-Venus (yellow) and anti-Dlg staining (cyan) reveal disruption of cellular organization in transformed follicle epithelial cells. (F, G) OC tumors fail to differentiate into the mature Hnt-positive (red), FasIII-negative (cyan) follicles seen in control. (H) Flies carrying OCs have greatly reduced survival compared to control, non-tumor-bearing flies. Tumor-bearing flies indicates substantial fluid accumulation in their hemocoel as indicated by macroscopic abdomen distention (I, J) and quantification of water weight (K). (L, M) Lipid staining (red) shows decreased fat tissue in OC flies. (N) Flies carrying OCs exhibit increased permeability of the blood-brain barrier. Scale Bars = 50μm (B,C,F,G), 25μm (D,E), 1mm (I,J), 100μm (L,M); Error bars = S.D.; *p < .05, ***p < .0005

Defining characteristics shared by mammalian and fly carcinomas include overproliferation, loss of cell polarity, and defective differentiation ^7,13^. Phosphohistone H3 staining revealed elevated mitotic rates in OC cells (**Figure 1B, C**). Epithelial organization was strongly disrupted, as was the localization of apically and basolaterally polarized proteins (**Figure 1D, E, S2**). Tumor cells failed to upregulate Hindsight (Hnt), a marker of mature differentiated follicle cells, and exhibited prolonged expression of the early follicle cell marker FasIII (**Figure 1F, G**) ^14^. Thus, fly OC cells are transformed into malignant epithelial tumors.

An additional characteristic of malignant tumors is their potential to kill hosts. Importantly, OC-bearing flies show dramatically accelerated mortality compared to controls, with median survival reduced by ∼50% (**Figure 1H**). GFP labeling of oncogene-expressing cells revealed no dissemination, suggesting a lack of metastasis (**Figure S1**). Since female flies do not require ovaries to live, lethality appears to arise from systemic effects of the tumor on the host, often called paraneoplasias. Similar to other adult tumor models^15–18^, OC-bearing flies exhibit bloating resulting from the accumulation of fluid in the hemocoel (**Figure 1I - K)**. Over the course of tumorigenesis, OCs also induce changes in fat body lipid storage (**Figure 1L, M**) ^15–17,19^. Finally, tumor-bearing hosts displayed breakdown of the blood-brain barrier (BBB) (**Figure 1N**) ^20^. The OC model therefore readily recapitulates previously documented *Drosophila* paraneoplasias.

To better understand OC progression and its impact on host survival, we developed a system to grade OC tumors based on distinct morphological characteristics (**Figure S2**). Assessing grade was more appropriate than mass because OC tumors, particularly at early grades, contain non-transformed germline cells that are polyploid and large. We defined grade 1 tumors as exhibiting a loss of epithelial polarity and overproliferation, most evident at the poles of individual follicles. While grade 1 tumors do not disrupt follicle organization and transformed cells remain *in situ*, grade 2 tumors show fusion between individual follicles as well as germline cell death. At grade 3, the muscle sheath and its associated basement membrane breaks down, allowing fusion between neighboring ovarioles. Further disruption of ovariole morphology resulting from organ-wide basement membrane breakdown characterizes grade 4 tumors. Using this system, we found that tumor grade reliably increased with time after induction, and that most dying animals displayed grade 3 or 4 tumors regardless of that individual’s time of death (**Figure S3**). The accelerated host mortality driven by tumor progression in a non-essential organ in the absence of metastasis place the OC model in a unique position to study tumor-host interactions.

### The OC transcriptome reveals candidate paraneoplasia mediators

To identify factors regulating these interactions, we performed bulk transcriptome analysis comparing wildtype (WT), pre-vitellogenic follicle cells to OC follicle cells 20 days after tumor induction (ATI). Expression of *matrix metalloproteinase 1* and *puckered* were increased, indicating high levels of JNK activity, as were *midline fasciclin* and *SOCS36E*, suggesting increased STAT activity (**Figure 2A**) ^21,22^. Fluorescent reporters validated the increased activity of both pathways (**Figure S4**), similar to other *Drosophila* epithelial tumors. *GstS1* and *Zfh*, markers of follicle stem cells and early prefollicle cells respectively^14^, were upregulated alongside downregulation of *Hnt* which marks more mature follicle cells, suggesting that some OC follicle cells retain a progenitor-like identity (**Figure 2B**).

**Figure 2.**
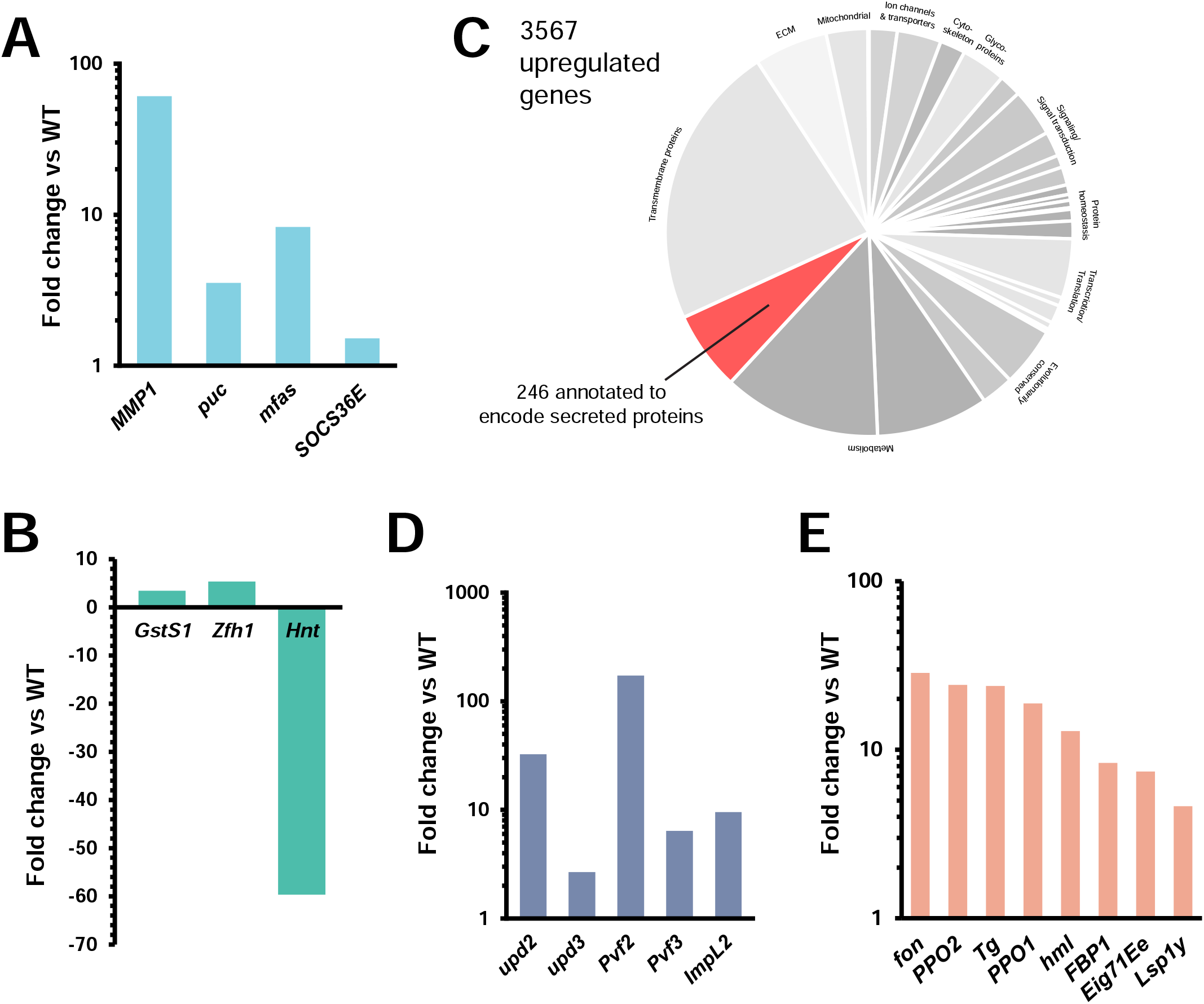
OCs differentially express clotting factors and regulators, among a number of other secreted proteins. (A) OCs exhibit heightened transcription of JNK pathway targets *MMP1* (60.97X) and *puc* (3.53X), as well as STAT pathway targets *mfas* (8.28X). (B) Increased proportion of OC tumor cells in early differentiation is indicated by enhanced transcript levels of *GstS1* (3.45X) and *Zfh1* (5.35X), and the failure to upregulate *hnt* (-59.71X). (C) 146 of 3567 transcripts upregulated in OC tumors vs control follicles are predicted to encode secreted proteins. (D) Like other *Drosophila* tumors, OCs upregulated *upd2* (32.58X) and *upd3* (2.68X), *Pvf2* (173.90X), *Pvf3* (6.41X), and *ImpL2* (9.54X). (E) Expression of many factors associated with hemolymph clotting was increased in OCs compared to wild-type follicle cells: *fon* (28.58X), *hml* (12.89X), *Lsp1γ* (4.62X), *Tg* (23.92X), *PPO1* (18.84X), *PPO2* (24.22X), *FBP1* (8.35X) and *Eig71Ee* (7.44X).

Since tumor-released peptides are likely mediators of paraneoplasias, we then focused on putative secreted proteins upregulated in the OC transcriptome. Of 3567 genes showing at least 2-fold increase overall, 246 are predicted to encode secreted factors (**Figure 2C**). Several of these encode known signals overproduced by adult gut tumors as well as larval disc tumors (reviewed in ref. 10), including the IL-6 like Unpaireds, PDGF/VEGF-related factors, and the Insulin Growth Factor Binding Protein-like ImpL2 (**Figure 2D**).

### Pro-clotting factors are upregulated in OC cells

Strikingly, among the upregulated genes predicted to encode secreted factors, we noted multiple genes that participate in the Drosophila clotting cascade (**Figure 2E**). As in mammals, flies have an essential circulatory fluid whose loss following wounding must be prevented. In both mammals and flies, this happens through clotting, in which soluble factors are polymerized and crosslinked to restore hemostasis ^23–25^. Genes whose products are found in larvae to form the initial ‘soft clot’ were strongly overexpressed in tumor cells, including *fondue (fon), hemolectin, Lsp1y, fat body protein 1, and Eig71Ee*, which can be considered functionally analogous to human fibrin ^26–28^ Some of these gene products are substates of the fly *Transglutaminase (Tg)*, which is homologous to mammalian clotting factor XIIIa ^29^, and is also upregulated in OCs. The clotting pathway in flies includes an additional insect-specific reaction, called the melanization cascade^30^. This reaction is responsible for clot maturation to form a ‘hard clot’ and is regulated by the activity of pro-phenoloxidases (PPOs), which are released by specialized blood cells called crystal cells (CCs) following injury ^30–33^. Transcript levels of *PPO1* and *PPO2* were elevated in OC cells as well. Taken together, these data raise the possibility that fly tumors may interface with the host clotting cascade.

### OC tumors induce coagulopathies in hosts

We therefore asked whether coagulation is altered by OC tumors. We first measured the capacity of hemolymph from tumor-bearing adult flies to form soft clots. Adapting an assay previously used for larvae ^34,35^, we incubated inert Dynabeads with extracted hemolymph *ex vivo*. Hemolymph from control adults at day 5 and 10 induced clumping of beads, albeit less strongly than seen in larvae (**Figure S5**). By contrast, hemolymph from OC flies 5-10 days ATI readily generated large, macroscopic bead aggregates, indicating that these animals are in a hypercoagulable state (**Figure 3A**). Aggregation activity subsequently decreases at days 15 and 20 ATI, demonstrating that the hypercoagulable state triggered by tumors is transient.

**Fig. 3:**
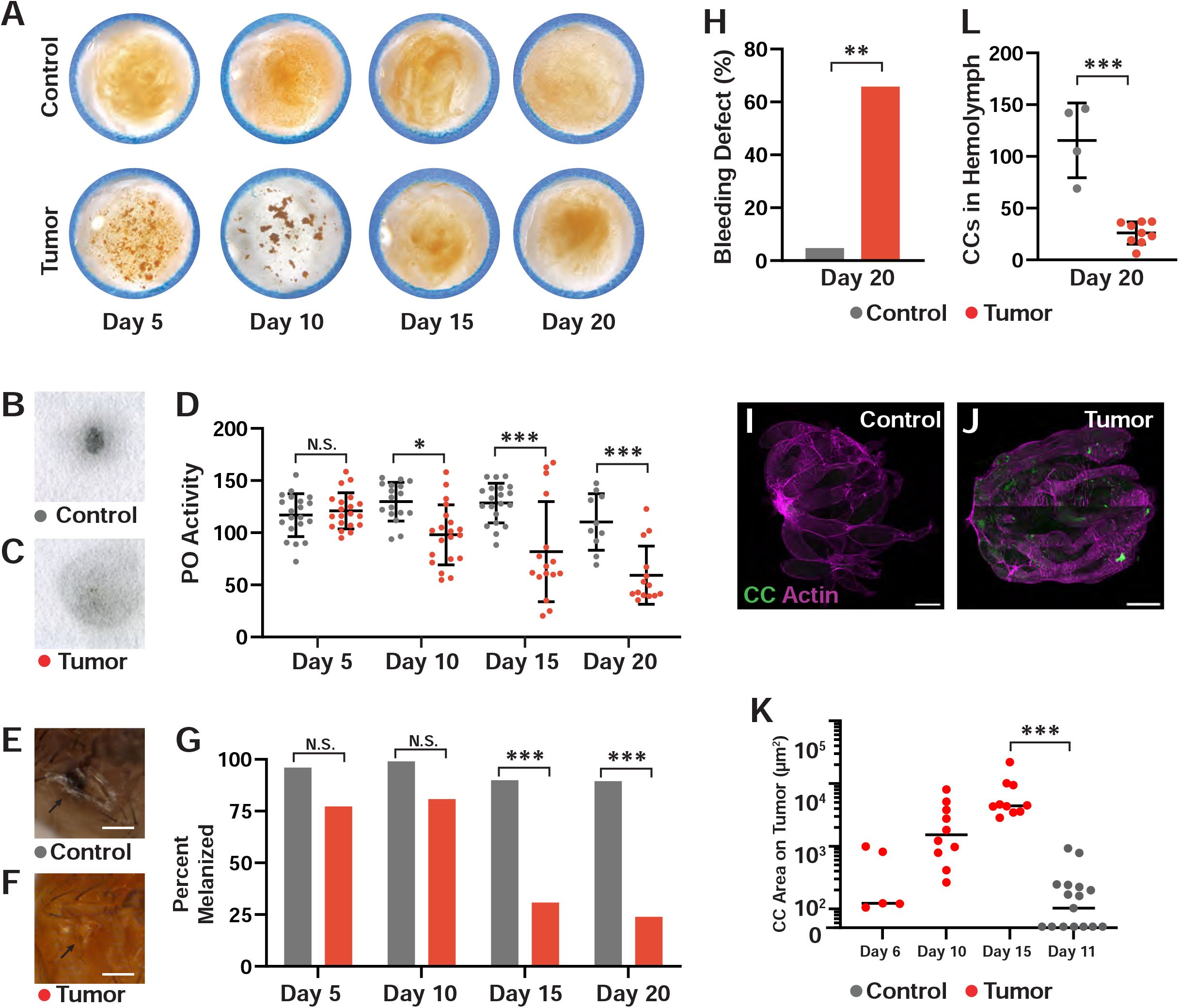
*Drosophila* ovarian tumors induce multiple defects in clotting. (A) Bead aggregation assay shows that soft clotting activity of control adult hemolymph (top) is dramatically increased in OC adult hemolymph (bottom) through day 10 ATI. Images are representative of multiple experiments and do not come from a single time course. PO activity in control (B) and OC flies (C) measured by L-DOPA blot reactivity. (D) Quantification of PO activity reveals decreases in OC flies following 10 days ATI. Thoracic wound response in control (E) and OC flies (F). (G) Percentage of flies able to melanize wounds reveals strong differences on days 15 and 20 ATI. (H) Hypocoagulation of OC flies, demonstrated by ability to measure hemolymph on throracic wounds after 2 hours. Crystal cells are rarely found on control ovaries (I) and accumulate on day 15 ovarian tumors (J). (K) Crystal cell area on tumors increases over time. (L) Crystal cell counts in hemolymph decrease in tumor-bearing flies at day 20 ATI. Scale Bars = 250μm; Error bars = S.D.; *p < .05, ***p < .0005

We then tested the ability of tumor-bearing hosts to create hard, melanized clots. We used two assays: first, measuring PO activity levels of hemolymph *ex vivo* on a colorimetric substrate ^33,36^, and second, by observing melanization of cuticular wounds *in vivo* ^37^. At day 15 ATI, PO activity was decreased in tumor-bearing flies in comparison to control (**Figure 3B, C**), and this decrease became more severe at day 20 (**Figure 3D)**. Tumor-bearing flies at days 15 and 20 ATI also showed a strong failure to melanize thoracic wounds compared to control flies (**Figure 3E - G**). Indeed, two hours after wounding, bleeding was staunched in control flies, but hemolymph remained on the wounds of tumor-bearing flies 20 days ATI (**Figure 3H**). The defects in hard clotting along with wound-healing are consistent with a hypocoagulable state that follows the hypercoagulable phase.

### Tumor-induced clotting creates a sink for clot cascade components

Given that hard clotting deficiencies temporally follow a period of increased soft clotting activity, we hypothesized that, as with some human paraneoplastic coagulopathies ^3^, fly tumors overstimulate the clotting system, leading to an exhaustion of clot components within the host. We tested whether OC-induced ectopic clots might act as a sink for limiting factors in the clotting cascade. A good candidate for such a factor are CCs, which lyse to release PPOs following activation and are produced only in the larvae but persist into adulthood ^38^. CCs, labelled by BCF2::GFP, were rarely detected on control ovaries (**Figure 3I, K**). However, at day 10, near the end of the hypercoagulable phase, multiple GFP-expressing cells were seen on OC tissue, and this became significantly higher than control at day 15 (**Figure 3J, K**). Lysis of these non-renewing CCs would be consistent with the observed depletion of hemolymph PO activity, contributing to clotting failure **(Figure 3B - G**). Indeed, counts of CCs in OC hosts revealed a decrease 15 days ATI compared to control (**Figure 3L)**. These results are consistent with the OC model causing a coagulopathy that consumes essential components of the clotting cascade.

### Coagulopathies driven by tumor upregulation of clotting components regulate host mortality

Finally, we asked whether coagulopathy plays a role in the premature lethality of OC flies. We tested this by using RNAi to deplete Fon, which is overproduced >25 fold in the tumor and is required for clotting in larvae ^39^. Remarkably, knockdown of Fon in OC tumors extended median survival by ∼38% (**Figure 4K**). This extension was robust compared to carefully matched control constructs, and is of a magnitude comparable to the lifespan extension of tumor-bearing hosts seen when blood-brain barrier disruption is prevented in a different tumor model ^20^. Increased lifespan correlated with reduced coagulopathy, since knocking down Fon attenuated early hypercoagulation as well as late hypocoagulation phenotypes. Quantitation revealed that hemolymph of flies bearing OC tumors depleted of Fon at day 5 ATI aggregated beads to a much lower extent (**Figure 4A - C**). At day 20 ATI these flies showed higher hemolymph PO activity and increased ability to melanize wounds compared to OC flies alone (**Figure 4D - J**). Thus Fon upregulation in the tumor regulates systemic coagulopathy and paraneoplastic lethality in the host.

**Fig. 4:**
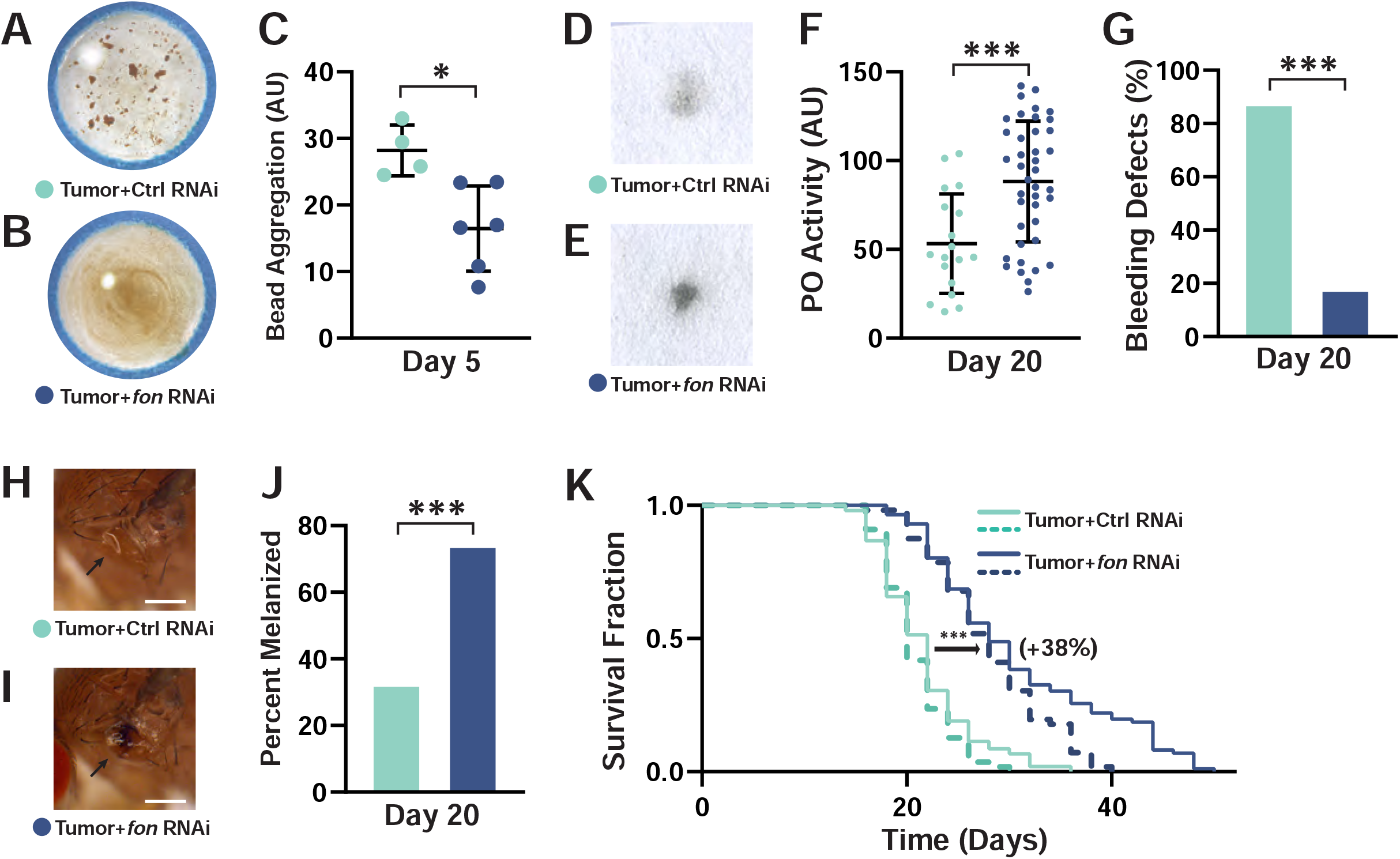
Tumor-secreted Fondue drives coagulopathies and early mortality. Compared to control RNAi depletion in OC cells (A), Fon depletion in OC cells (B) rescues the strong hypercoagulation displayed by tumor-bearing flies, revealed by the bead aggregation assay quantitated in (C). Compared to control RNAi depletion in OC cells (D), Fon depletion in OC cells (E) also ameliorates the loss of PO activity, quantitated in (F), and defects in bleeding (G) and wound melanization (H-J). (K) Early mortality induced by OC tumors is significantly reduced upon Fon depletion. Solid and dashed lines represent different replicates of the experiment. Scale Bars = 250μm; Error bars = S.D.; *p < .05, ***p < .0005

One explanation for coagulopathy accelerating TBH mortality could be an autonomous effect on tumor progression. Yet quantitation of OC grade revealed no change when Fon was depleted (**Figure S6A, B**). A second possibility is that coagulopathy might reduce lifespan through enhancing BBB permeability, which has been recently shown to contribute to tumor-associated lethality ^20^. BBB breakdown was present in OC flies even upon depletion of Fon (**Figure S6C**). We investigated a third possibility –that coagulopathy causes an elevation of ROS levels through CC release of PPOs. ROS has well-documented deleterious effects on lifespan, but also has been suggested to have cytoprotective effects ^40,41^. We found that OC flies heterozygous for the *Bc* mutation, which causes premature rupture of many CCs during the larval stage, show no changes in lifespan, nor do OC flies carrying an engineered dominant PPO1 allele with similar effects on CCs (**Figure S7A, B**) ^42,43^. Furthermore, treatment with antioxidant or pro-oxidant compounds under different regimes did not significantly change OC mortality (**Figure S7C, D**). Finally, dying OC flies did not show increased intestinal permeability as detected by the ‘smurf’ assay (**Figure S7E**) ^44^. Overall, these data suggest that coagulopathy contributes to tumor-driven death via a currently unknown paraneoplastic mechanism.

Our data reveal a remarkable and unexpected parallel between cancer patients and tumor-bearing flies: both can show widespread clotting defects that contribute to lethality. Systemic coagulopathies are common in cancer patients and have been studied since Trousseau’s syndrome was described in 1865 ^3,4^. One coagulopathogenic mechanism of many human tumors is abnormal upregulation of Tissue Factor, which can trigger the clotting cascade ^45^. Although insects do not have a similar single coagulant-initiating factor, the broad cocktail of clot-regulating proteins secreted by OC tissue suggests an analogous response in fly tumors. Indeed, to our knowledge, the tumor data shown here reveal the first hypercoagulative phenotype documented in the fly. This adds to compelling evidence that the similar physiological reaction of the host to wounds and tumors has a quite ancient origin with a deeper level of conservation than previously appreciated.

Clotting complications are the second leading cause of death for cancer patients, primarily through venous thromboembolisms (VTEs) which are detected in 10-15% of all malignancies ^4^. However, some evidence suggests that coagulopathy may also impact mortality through non-thrombotic pathways as well. Elevated markers of clotting factors in circulation are strongly correlated with poor survival, yet thrombosis is often not observed in patients ^5,6^. There is also some evidence that prophylactic treatment with anticoagulants can improve cancer patient outcomes beyond prevention of thrombosis ^46^. Thus, there may be unappreciated mechanisms through which altered clotting behavior contributes to morbidity and mortality. The fly, with its open circulatory system, is unlikely to be dying from tumor-induced thrombosis, and thus may be used as a discovery system for potential alternative mechanisms. In this work, we have not determined the exact reason why coagulopathy promotes the death of tumor-bearing flies, but our data argue against several *prima facie* feasible possibilities.

The initial hypercoagulative phenotype of OC flies followed by hypocoagulation echoes features of patient conditions such as Disseminated Intravascular Coagulopathy (DIC). DIC is considered a consumptive coagulopathy, where ectopic activation of pro-coagulation pathways can paradoxically lead to excessive bleeding through local depletion of hemostatic components^2^. Our data point to fly CCs, which lyse to provide PO clot-hardening activity, as one consumed regulator, but hint that the dramatic hypercoagulation, rather than hypocoagulation, of tumor bearing flies may drive the detrimental consequences for mortality. Future studies will explore how this simple fly system reacts to the tumor-induced danger response, perhaps by triggering inflammatory, immune, or metabolic responses that have a negative impact on host survival.

## Acknowledgements

We acknowledge Ulrich Theopold for sharing Drosophila clotting expertise, and Yuejiang Liu for technical assistance. Iswar Hariharan, Lin He, Daniel Nomura and Shaina Carroll provided many helpful discussions. Reagents were generously gifted by Bruno LeMaitre, Robert Schulz, Katja Brueckner, Elisabeth Knust and the community resources by the Vienna Drosophila Resource Center and the Bloomington Drosophila Stock Center. This work has been supported by NIH grants GM130388 and GM090150 to DB, a University of California Cancer Research Coordinating Fellowship to TH, and a Mark Foundation-Damon Runyon Fellowship 2400-20 to KO.

## MATERIALS AND METHODS

### Fly husbandry and stocks

Flies were maintained on cornmeal, molasses, and yeast food at 21°C in wide vials unless otherwise noted. A complete list of stocks used is given in **Table S1**. The *fon* RNAi construct was validated by confirming that it can recapitulate previously described larval soft clotting and pupariation defects ^39^.

### Ovarian tumor induction and lifespan assays

After eclosion, adult flies were kept at 21°C on food with yeast powder for two days. No more than 25 flies were kept in each wide vial, a density that was previously determined did not impact longevity. Flies were then put on new food and shifted to 29°C to initiate tumor induction. Food was changed every two days and the number of dead flies was counted. For each lifespan assay, at least 50 flies were used for each sample group and all assays were repeated twice, with control and experimental groups run in parallel. For RNAi lines used in lifespan assays, the lines were backcrossed at least four generations to *w*^*1118*^ (BL#5905) to minimize variation in genetic background between stocks.

### Tumor transcriptome sequencing and data analysis

Twenty days after shifting to restrictive temperature, around two hundred *tj-GAL4 tubulinGal80ts* and *tj-GAL4 tubulinGa80ts; UAS-aPKCΔN UAS-RasV12* ovaries were dissected in Schneider’s *Drosophila* Media (Life Technologies, 21720024) on ice for each biological replicate. At least two biological replicates were sequenced per genotype. Follicles ∼stage 6 and older of control ovaries were removed by cutting the distal regions. Ovaries were washed three times in DPBS (Dulbecco’s Phosphate-buffered saline; Life Technologies, 14190144) followed by 10 minute incubation in 10mg/mL *Bacillus licheniformis* protease (Sigma, P5380) prepared in DPBS at 6°C while vigorously agitating. Protease activity was quenched by adding a half volume of 10% fetal bovine serum (FBS) in Schneider’s *Drosophila* Media. Cell suspension was then filtered using a 50μm filter (Sysmex Partec, 04-004-2327) to remove germ cells, and the filtered suspension was subsequently spun down for 7 minutes at 1000g. RNA of pelleted cells was isolated using RNeasy mini kit (QIAGEN, 74104), and RNA quantity and quality was analyzed by the Functional Genomics Laboratory at the California Institute for Quantitative Biosciences at UC Berkeley (QB3-Berkeley).

Libraries were sequenced by 50bp single-end reads on HiSeq4000 platform (Illumina, San Diego, CA), and sequencing was performed by the Vincent J. Coates Genomics Sequencing Lab at QB3-Berkeley. Sequences were aligned to the *Drosophila melanogaster* reference genome (version 6.26) using Kallisto under default parameters for single-end reads ^47^. Lowly expressed genes were removed prior to differential expression analysis using DESeq2 ^48^.

### Immunofluorescence

Ovarian tumors, thoracic muscle, brains and intestines were dissected in PBS and fixed in 4% PFA-PBS for one hour at room temperature without agitation. Samples were washed three times with PBS-TX (0.1% Triton-X in 1X PBS) before proceeding to dye treatment. Samples were blocked in 2% BSA or 4% NGS/1% BSA blocking solution for 30-60 minutes. Primary antibodies were incubated overnight at 4°C and used at the following concentrations: anti-Dlg (1:100), anti-phospho-histone H3 (1:250), and anti-Hindsight (1:100). Samples were washed three times with PBS-TX and incubated with AlexaFluor-conjugated secondary antibodies for 1 hour at room temperature. Tissues were again washed three times with PBS-TX prior to additional chemical staining. To stain actin, fixed tissues were incubated for 30-60 minutes at room temperature with Rhodamine-Phalloidin at a concentration of 1:500. To stain nuclei, a 1:1000 DAPI solution was applied for 10 minutes. After all incubations, samples were washed three times with PBS and incubated in the final wash for at least 20 minutes. Samples were mounted in Diamond Antifade Mountant before imaging.

### Visualizing fat body wasting

Dorsal cuticles were dissected from adult flies in PBS and fixed in 4% PFA for 30 minutes. Samples were washed once in PBS-TX and three times in PBS. Dorsal cuticles were incubated in Nile Red staining solution (0.5 μg/mL in 1X PBS) for 10 minutes followed by three washes in PBS. Samples were mounted in Diamond Antifade Mountant and stored at 4°C. Images were taken at multiple focal planes on Zeiss AxioImager M2 using Zeiss Zen (blue edition 2.3) pro imaging software, and multifocus images were created using Helicon Focus 7.

### PO activity measurements

We used a modified protocol from ref. 36. To measure PO activity of hemolymph *ex vivo*, individual flies were bled onto small squares of Whatman filter paper (1cm^2^; Whatman 1001 – Grade 1). Immediately following each bleeding, 20μL of 20mM L-DOPA in PBS was applied to each blot. Samples were then covered to prevent evaporation and incubated for 30 minutes at 25°C. The blots were rapidly dried by heating in a microwave for 10 seconds and then allowed to completely air-dry for 30 minutes. Following drying, each blot was sealed in clear Scotch tape and scanned using an Epson Perfection 4490 Photo Scanner. Intensity of each blot was quantified in FIJI, and the mean background intensity was subtracted from all samples.

### Thoracic wound healing assay

Flies were anesthetized with CO_2_, and one side of the thorax was punctured using a blunt 0.005 inch diameter tungsten needle (Ted Pella #27-11). Flies were then returned to vials, and two hours later counted for melanized thoracic wounds using a dissecting microscope. To determine wound-healing failure, a glass micropipette tip (World Precision Instruments #TIP30TW1) was gently brought into contact with the thoracic wound two hours after wounding, and assessed for whether hemolymph was drawn up by capillary action.

### Bead aggregation assay

We used a modified protocol from ref. 35. Dynabeads were washed with 10X PBS twice and blocked in a 0.1% BSA-PBS solution overnight on a rotator at 4°C. The beads were washed three times with 0.1X PBS and reconstituted in Ringer’s-PTU (130mM NaCl, 5mM KCl, 1.5mM CaCl_2_ x 2H_2_O, 2mM Na_2_HPO_4_, 0.37mM KH_2_PO_4_, 0.01% PTU) buffer at a final concentration of 50%. The blocked beads were stored up to two weeks on a rotator at 4°C. To extract hemolymph, we used a modified protocol from ref. 49. A Qiagen spin column was disassembled and the filter paper removed. All other components were rinsed in MilliQ purified water. The column was reassembled without the filter and centrifuged at 13.2 × 1000g for 10 minutes at room temperature. The tube that the column was nested in was replaced with a fresh 1.5mL Eppendorf tube. Flies were anesthetized and shallowly cut across the dorsal thorax with a 33G needle. The flies were transferred to the modified column stored on ice. All flies were wounded in less than 10 minutes to avoid loss of clotting activity. The column was spun at 5000g for 5min at 4°C. 2μL of hemolymph was combined with 2uL of blocked beads in a well of a 15-well glass slide by pipetting up and down ten times. The slide was incubated in a humid chamber at 25°C for 30minutes. Bead aggregates were revealed by swirling the solution with a 10μL micropipette tip for 30 seconds. The well was photographed with an iPhone 12 camera mounted on a dissecting microscope. To measure bead aggregation, we used the Bernsen Adaptive Local Thresholding Method available in FIJI to generate a binary image. A polygon was drawn around the liquid droplet and the percent area unoccupied by beads (i.e. the white area) was measured. The final measurement was generated after subtracting the average white area percent measured across five wells of control reactions with 2μL of Ringer’s-PTU buffer instead of hemolymph.

### Confocal microscopy

Fixed samples were imaged on the Zeiss LSM700 Scanning Confocal Microscope with a Plan-APOCHROMAT 20x/0,8 objective. The microscope was controlled with Zeiss Zen 2010 imaging software. Images were processed and analyzed with FIJI software.

### Blood brain barrier permeability assay

Assays were done as described in ref. 20. Approximately 100nL of 25 mg/ml 10,000 MW TR-dextran was injected in the abdomen of adult females using a fine glass needle. If tumor-bearing hosts excreted hemolymph upon injection due to bloating, excess hemolymph was removed. 15 hours after injection, flies were fixed in 4% PFA-PBS for 80 minutes and washed in 0.1% PBS-TX before brain dissection. Brains were imaged on the same day of fixation to minimize diffusion of dextran. Average intensity was measured in a cross-section at the center of the brain at two regions using FIJI software.

### Drug Treatments

Paraquat was used at either 2mM or 5mM. N-Acetyl Cysteine (NAC) was administered at 20mM. All drugs were dissolved in MilliQ-filtered water. 0.74g of 4-24 Carolina Instant Blue Food was mixed with 2mL of water or drug cocktail. Drugged food was always made fresh the day of use. Paraquat treatment was started 6 days after tumor induction. After two days, flies were transferred to food reconstituted with water, and thereafter consistently alternated every two days between Paraquat food and water only food. This alternating treatment was done to avoid Paraquat-driven mortality ^50,51^. NAC treatment was begun 10 days after tumor induction to avoid potential delays in early tumorigenesis from the antioxidant treatment.

### Water weight measurements

Measurements were done as described in ref. 52. Briefly, four to five flies were placed into a 1.5mL centrifuge tube, and wet weight was measured on Mettler Toledo AG104 scale. Flies were then dried at 65°C overnight with the centrifuge tube opened to allow for evaporation. Dry weight was then measured. Wet-dry weight was obtained by subtracting the dry weight measurement from the wet weight measurement, and the result was divided by the number of flies in the respective tube.

### Smurfing Assay

The smurfing assay was carried out as described in ref. 44. In brief, flies were moved onto food containing 2.5% wt/vol Blue Dye No. 1. After 9 hours, the flies with dye present outside of the digestive tract were counted.

### Statistics

A one-way ANOVA test or Student’s T-test was used for all parametric data (crystal cell area, crystal cell counts, PO activity, bead aggregation, water weight and BBB permeability). The Kruskal-Wallis test or Fisher’s exact test was used for all non-parametric data (wound melanization, tumor grading, bleeding defects and smurf assay). The Log-Rank test was used to determine significant differences in survival curves. Graphpad Prism and Python were used to perform statistical testing.

## SUPPLEMENTAL FIGURE LEGENDS

**Figure S1:**
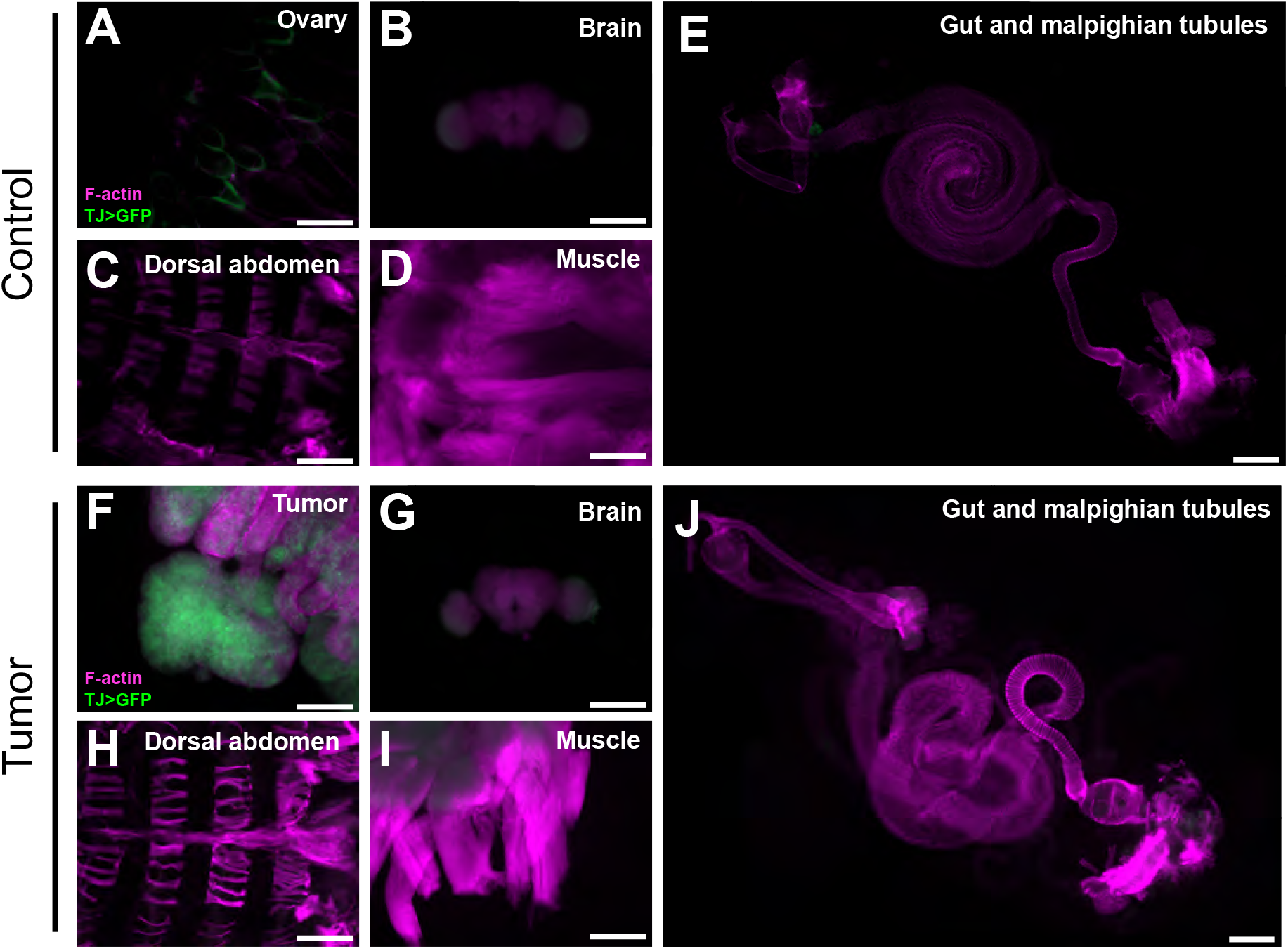
OC model lacks metastasis. Images from adult tissues of control (A-E) vs OC flies (F-J) expressing *UAS-NLS-GFP* 20 days ATI. Green fluorescence clearly marks cells in the control ovary (A) that show morphological transformation in OC flies (F). *tj-GAL4* shows some expression in the optic lobes but these structures do not obviously change upon coexpression of the OC oncogenes (B, G). No green cells are seen in muscle, abdomen, or digestive tract (C-E, H-J) with the exception of small GFP-labeled cells by the crop that are also present in OC flies. Scale bars = 250μm.

**Figure S2:**
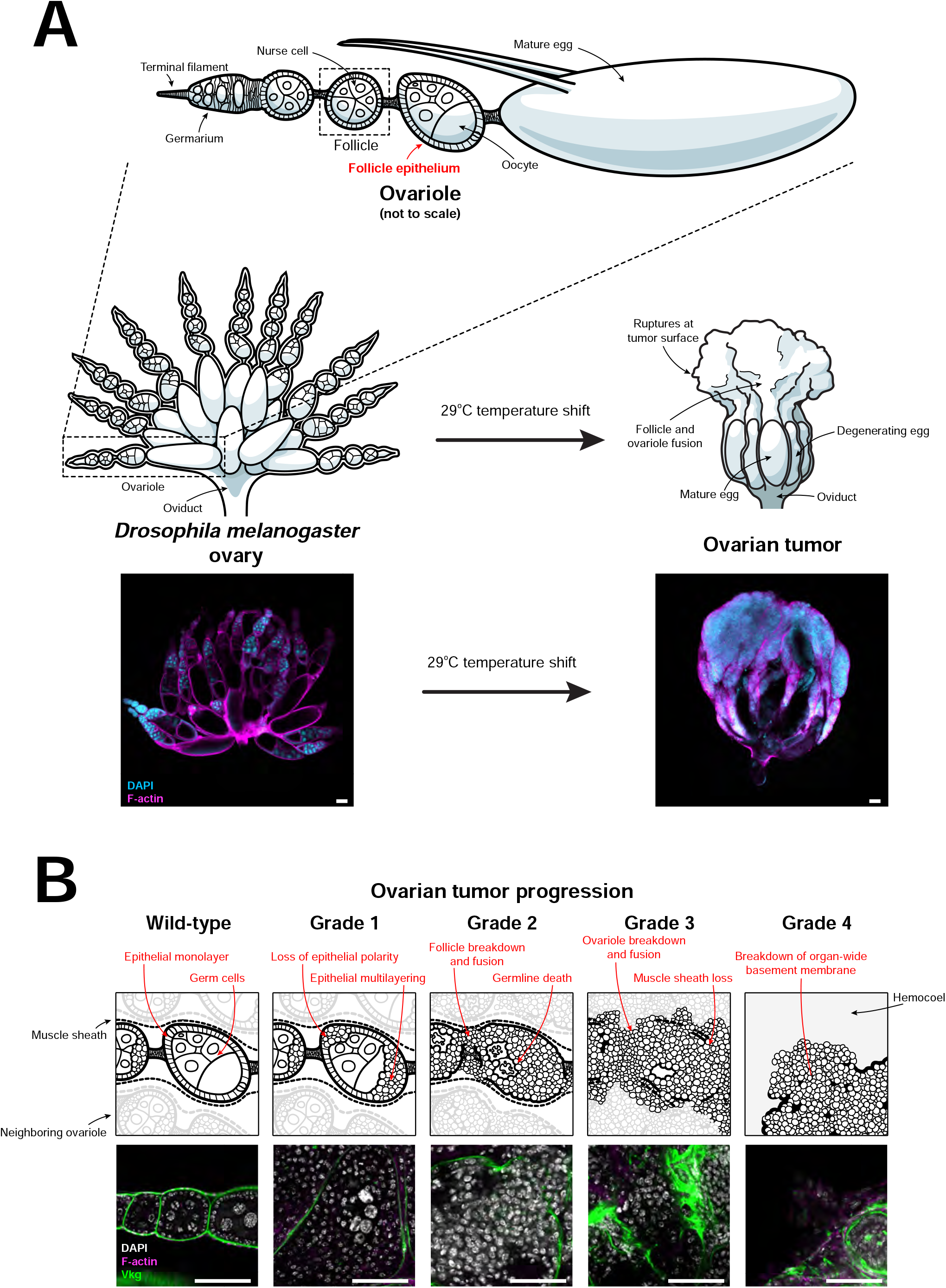
Adult Drosophila ovarian tumor characteristics and progression. (A) Schematic of the *Drosophila* ovarian carcinoma (OC) model illustrates the architecture of normal ovaries and the progression of malignant transformation of follicle epithelial cells. Representative confocal micrographs are shown below illustrations of ovaries. (B) Illustrations of distinguishing features of tumor grade stages 1-4 with corresponding representative micrographs below. Nuclear (white) and Actin (magenta) stains show dramatic changes in cellular and tissue organization. Vkg::GFP (green) reveals basement membrane breakdown. Scale bars = 100 μm (A), 50μm (B).

**Figure S3:**
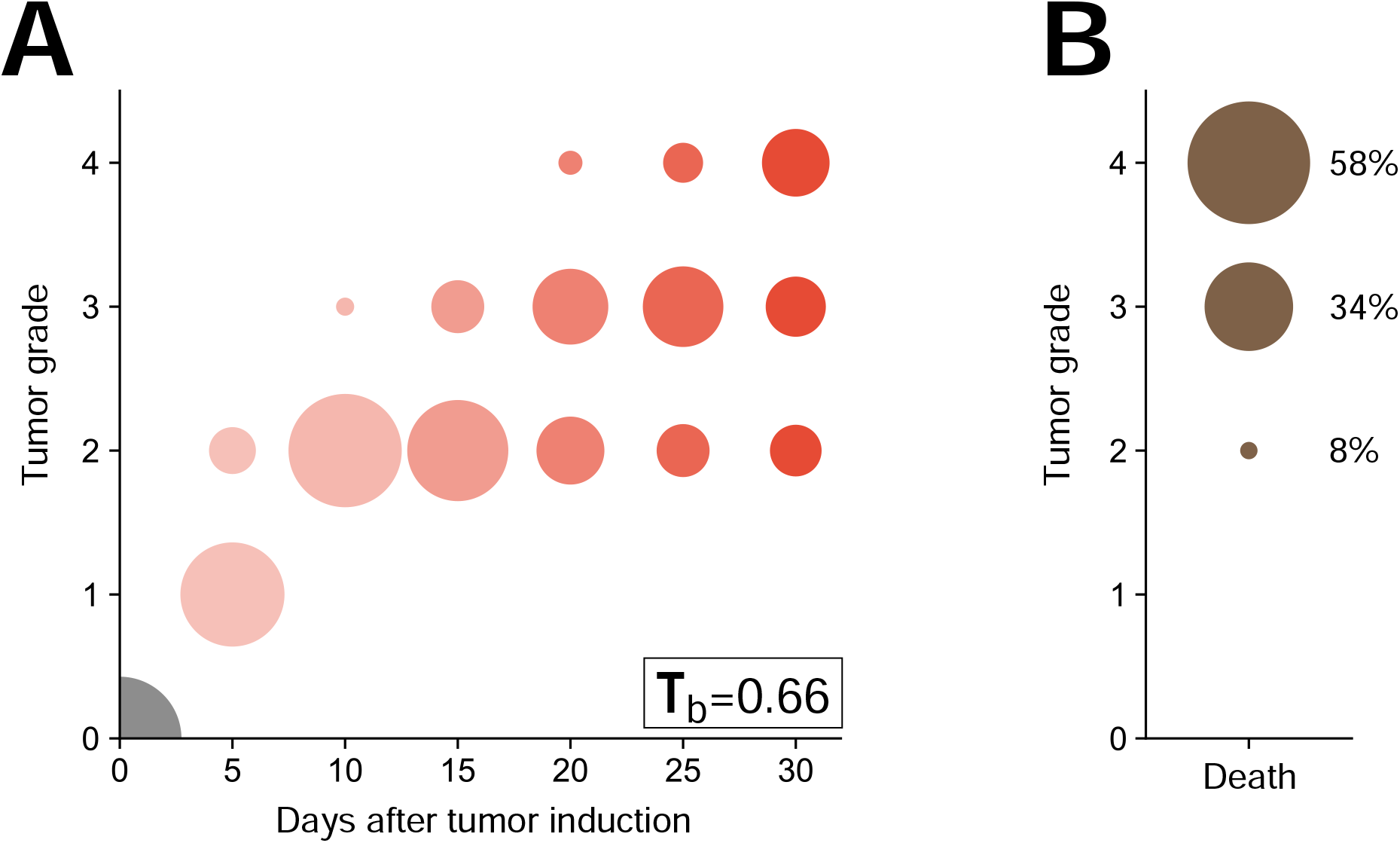
Higher tumor grade associated with lethality. (A) Tumor grade is well correlated with time after tumor induction, with a greater proportion of grade 3 and 4 tumors in later time points. Size of circle represents proportion of sample in each class. Correlation determined by calculating Kendall’s τ_b_ correlation coefficient; τ_b_=0.66. (B) The majority of flies necropsied following death between 10 and 28 days ATI had tumors of grades 3 or 4, indicating a strong association between tumor grade and its lethality.

**Figure S4:**
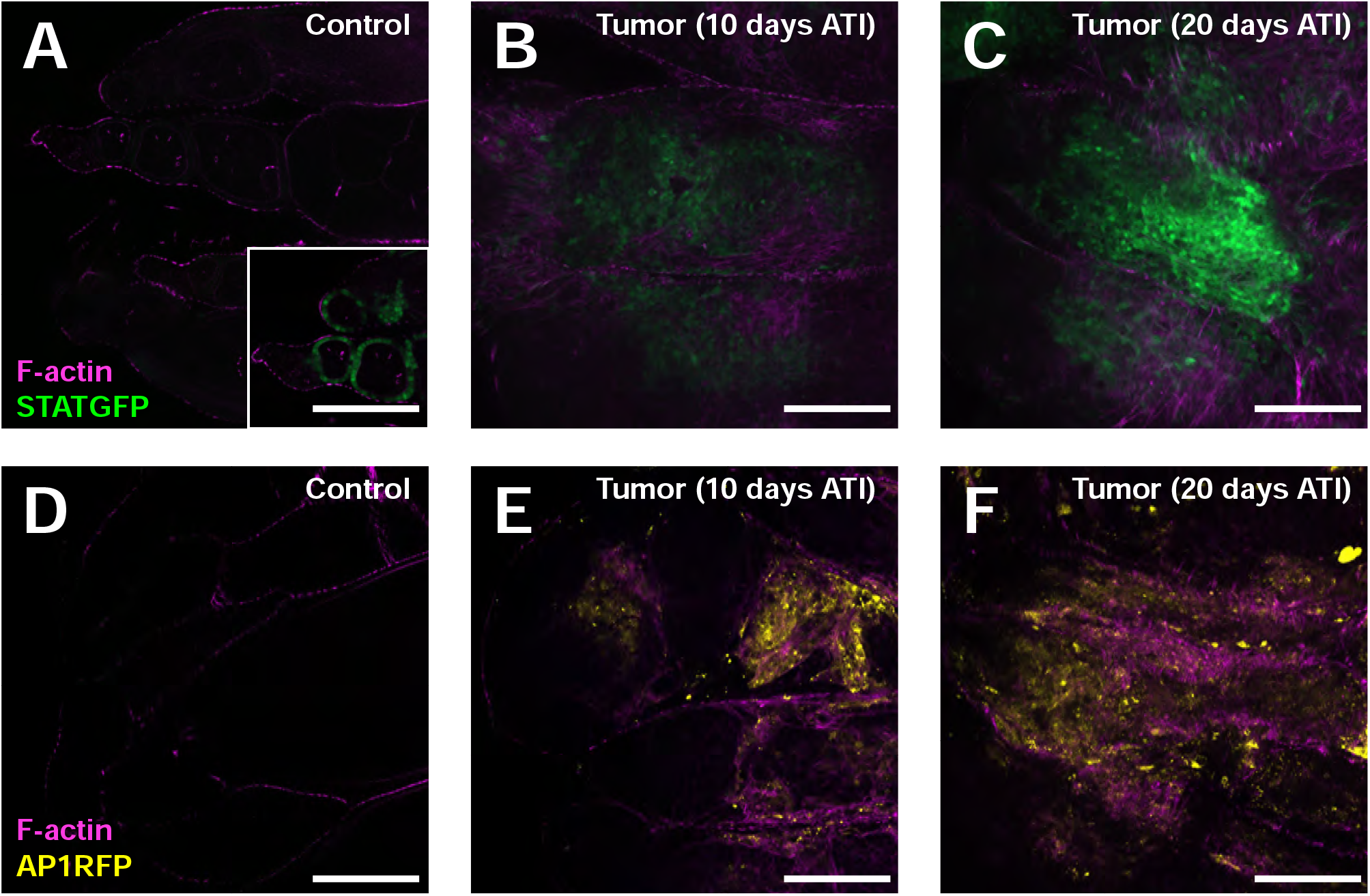
Jnk and Jak/STAT signaling in OC tumors. (A) Control ovaries show expected low-level Jak/STAT signaling (STATGFP activity reporter, green) in the follicle epithelium induced by Upd from polar cells. These signals are highlighted in inset using lower fluorescence threshold values. (B) Tumor at 10 days ATI shows elevated STAT signaling throughout (threshold values matched to main panel of A). (C) Tumor at 20 days ATI shows greatly increased STAT signaling (threshold values matched to main panel of A). (D) AP1 activity reporter (yellow) shows minimal Jnk signaling in control ovaries. (E) 10 days PTI and (F) 20 days ATI, Jnk signaling is strongly activated. Scale bars = 100μm.

**Figure S5:**
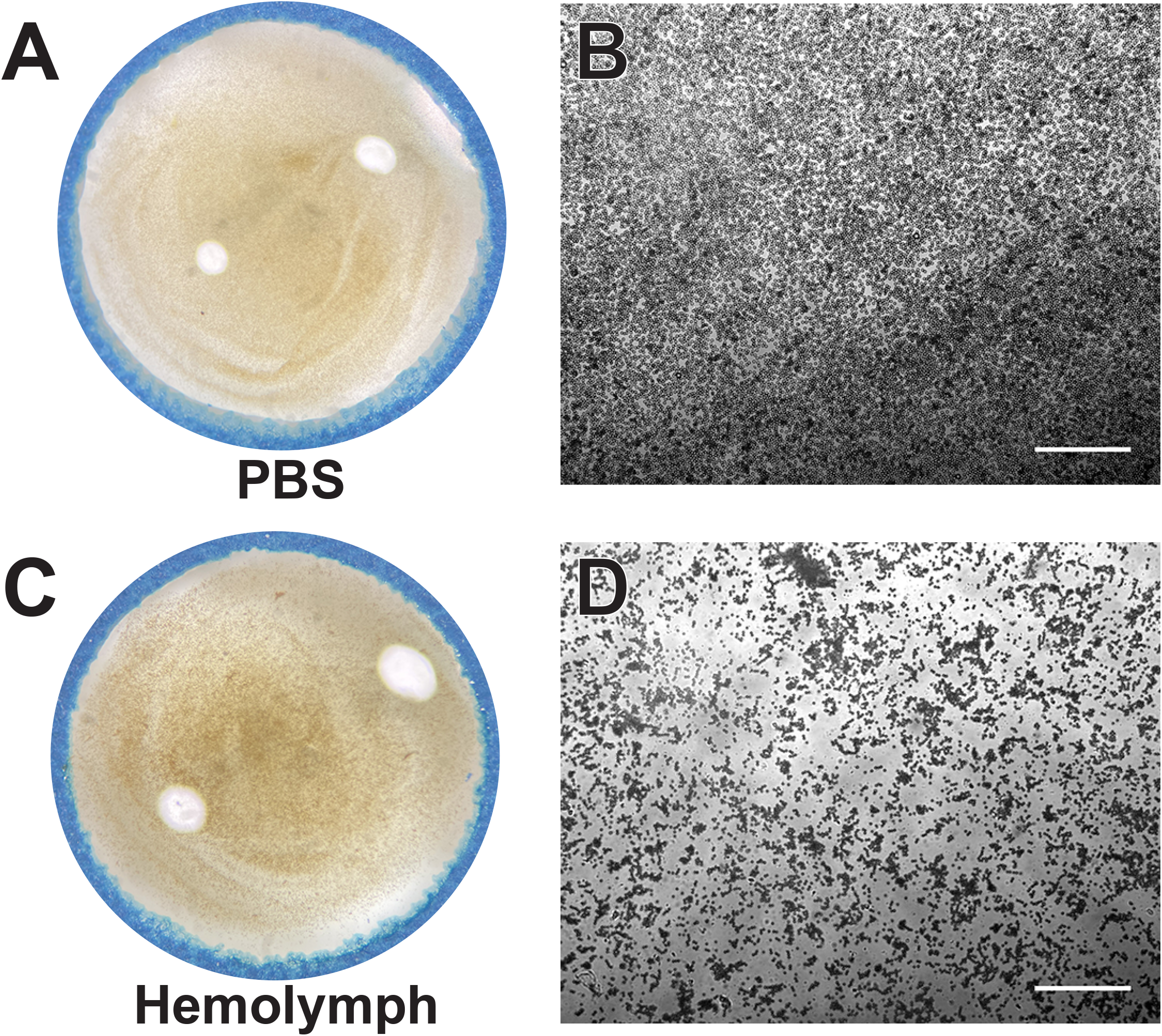
Clotting activity in adult fly hemolymph. 4μL of either PBS (A,B) or hemolymph from 14 day-old *w*^*1118*^ adult flies (C,D) mixed with bead solution reveals aggregation activity in adult *Drosophila*. The fine clot formed by adult fly hemolymph is more evident when beads are imaged microscopically rather than macroscopically. Scale bars = 100μm.

**Figure S6:**
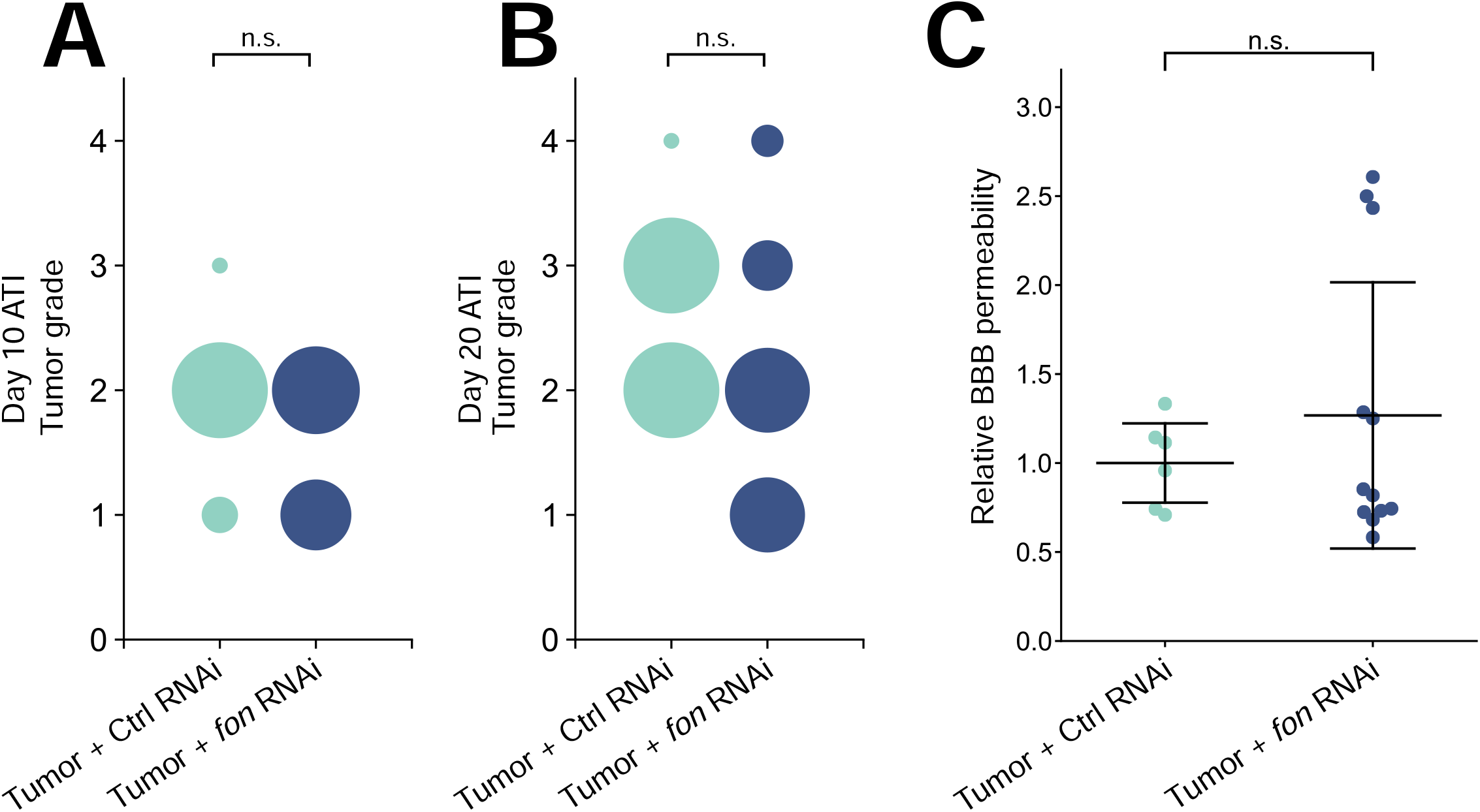
*Fon* depletion in OC tumors does not reduce tumor grade nor BBB permeability. Knockdown of *fon* via RNAi in OC tumors does not significantly change tumor grade at day 10 (A) or 20 ATI (B). Size of circle represents proportion of sample in each class. Knockdown of *fon* via RNAi in OC tumors does not significantly change BBB permeability at day 20 ATI relative to control RNAi (C).

**Figure S7:**
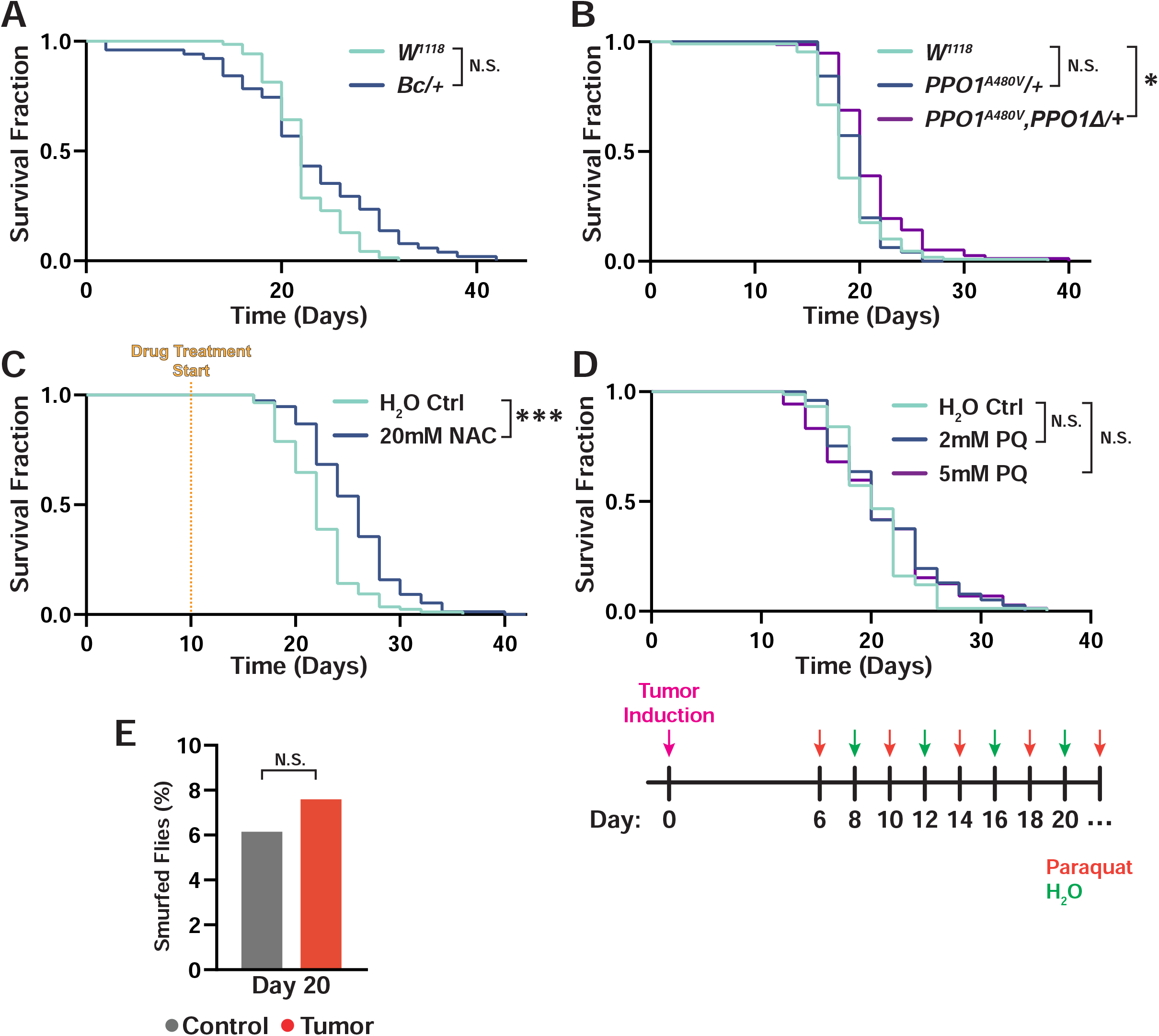
PO, ROS, and intestinal permeability do not account for early lethality in OC flies. (A) Tumor-bearing flies heterozygous for a dominant mutation in PPO1 (*Bc*/+) do not live longer than flies with normal PO levels. (B) Flies carrying a copy of an independently-generated dominant mutation in PPO1 (*PPO1*^*A480V*^/+) do not die significantly later than respective controls. The additional loss of a copy of PPO1 (*PPO1*^*A480V*^, *PPO1*^*Δ*^/+) only extends the mean survival by 1 day. (C) Reducing systemic ROS by treating tumor-bearing flies with the antioxidant N-acetyl Cysteine (NAC) only improved mean survival by 3 days. (D) Feeding tumor-bearing flies with paraquat (PQ), which increases ROS, did not significantly alter lifespan. Below, the PQ-feeding schedule with respect to tumor induction is shown. (E) OC flies at day 20 (median lifespan) do not show increased intestinal permeability, as assessed by the smurf assay. *p < .05, ***p < .0005.

**Table S1:** Reagents, chemicals, fly stocks, and programs used.

| REAGENT or RESOURCE | SOURCE | IDENTIFIER |
| --- | --- | --- |
| **Antibodies** |  |  |
| Mouse Anti-Dlg | Developmental Studies Hybridoma Bank | 4F3 |
| Anti-phospho-Histone H3 | Cell Signaling | 9701 |
| Mouse Anti-Hindsight | Developmental Studies Hybridoma Bank | 1G9 |
| **Chemicals, peptides, and recombinant proteins** |  |  |
| TRITC-Phalloidin | Sigma-Aldrich | P1951 |
| DAPI | ThermoFisher Scientific | D1306 |
| Nile Red | Sigma-Aldrich | N3013 |
| Paraformaldehyde 16% | Electron Microscopy Sciences | 15710 |
| Phosphate Buffered Solution | Sigma-Aldrich | P4417 |
| Triton-X | Sigma-Aldrich | T8787 |
| Phenylthiourea | Sigma-Aldrich | P7629 |
| L-DOPA | Sigma-Aldrich | D9628 |
| Dynabeads M-280 Tosylactivated | Invitrogen | 14203 |
| Formula 4-24 Instant Drosophila Medium | Carolina Biological Supply Company | 173210 |
| Bovine Serum Albumin | Sigma Aldrich | A3912 |
| 10 kDa Texas red-Dextran | ThermoFisher Scientific | D1863 |
| SlowFade Diamond Antifade Mount | Invitrogen | S36963 |
| TWEEN 20 | Sigma-Aldrich | P9416 |
| NaCl | Fisher Chemical | S271 |
| KCl | Fisher Chemical | P217 |
| CaCl<sub>2</sub> x 2H<sub>2</sub>O | Fisher Chemical | C79 |
| Na<sub>2</sub>HPO<sub>4</sub> | Fisher Chemical | S374 |
| KH<sub>2</sub>PO<sub>4</sub> | Fisher Chemical | P284 |
| Paraquat | Sigma-Aldrich | 36541 |
| N-Acetyl-L-Cysteine | Sigma-Aldrich | A7250 |
| Alexa Fluor Phalloidin 647 | Invitrogen | A22287 |
| Schneider's *Drosophila* Medium | Gibco | 21720-024 |
| Dulbecco's Phosphate-buffered saline | Gibco | 14190144 |
| Protease from *Bacillus licheniformis* | Sigma-Aldrich | P5380 |
| Fetal bovine serum | Cytiva Life Sciences | SH30070.03 |
| RNeasy mini kit | QIAGEN | 74104 |
| **Experimental models: Organisms/strains** |  |  |
| *w*<sup>1118</sup> | Bloomington Drosophila Stock Center | #5905 |
| *BcF2-GFP* | Gajewski et al 2007 | FBal0244649 |
| *BcF6-GFP* | Gajewski et al 2007 | FBal0244650 |
| *UAS-Fondue shRNA* | Vienna Drosophila Resource Center | 330748 |
| *UAS-GFP shRNA* | Vienna Drosophila Resource Center | #60201 |
| *10XSTAT-GFP* | Bloomington Drosophila Stock Center | #26198 |
| *AP1-RFP* | *Chatterjee & Bohmann 2012* | FBal0268835 |
| *Vkg::GFP* | Vienna Drosophila Resource Center | v318167 |
| *TJ-Gal4* | Kyoto Stock Center | #104055 |
| *UAS-aPKC<sup>ΔN</sup>* | Bloomington Drosophila Stock Center | #51673 |
| *UAS-Ras<sup>V12</sup>* | Bloomington Drosophila Stock Center | #4847 |
| *Fasciclin III::GFP trap line* | Bloomington Drosophila Stock Center | #59809 |
| *Tub-Gal80<sup>ts</sup>* |  |  |
| *UAS-SAS::Venus* | *Firmino et al 2013* | FBtp0084565 |
| *UAS-NLS-GFP* |  |  |
| Software and algorithms |  |  |
| FIJI |  | ImageJ.net |
| Zen 2010 | Zeiss |  |
| Zen (blue edition 2.3) pro | Zeiss |  |
| Kallisto Alignment Software | Pachter Lab | https://pachterlab.github.io/kallisto/ |
| R Statistical Software | The R Foundation | n/a |
| Graphpad | Prism | Ver 9.3.1 |
| Gene List Annotation for Drosophila | DRSC/TRiP Functional Genomics Resources | https://www.flyrnai.org/tools/glad/web/ |
| Python |  | Ver 3.9.7 |
| Other |  |  |
| 15-well slides | MP Biomedicals | 096041505 |
| Pre-pulled glass pipette -- 30 μm diameter | WPI | TIP30TW1 |
| 50 μm filter | Sysmex Partec | 04-004-2327 |
| Qiaprep 2.0 Spin Columns | QIAGEN | 27115 |

